# Analysis and comparison of genome editing using CRISPResso2

**DOI:** 10.1101/392217

**Authors:** Kendell Clement, Holly Rees, Matthew C. Canver, Jason M. Gehrke, Rick Farouni, Jonathan Y Hsu, Mitchel Cole, David R. Liu, J. Keith Joung, Daniel E. Bauer, Luca Pinello

**Affiliations:** Broad Institute of MIT and Harvard, Cambridge, Massachusetts 02141, USA; Molecular Pathology Unit, Center for Cancer Research, and Center for Computational and Integrative Biology, Massachusetts General Hospital, Charlestown, Massachusetts 02129, USA; Department of Pathology, Harvard Medical School, Boston, Massachusetts 02115, USA; Department of Chemistry and Chemical Biology, Harvard University, Cambridge, Massachusetts 02138, USA; Division of Hematology/Oncology, Boston Children’s Hospital; Department of Pediatric Oncology, Dana-Farber Cancer Institute, Boston, MA 02115, USA; Department of Pediatrics, Harvard Medical School, Boston, MA 02115, USA; Harvard Stem Cell Institute, Cambridge, MA 02138, USA

## Abstract

Genome editing technologies are rapidly evolving, and analysis of deep sequencing data from target or off-target regions is necessary for measuring editing efficiency and evaluating safety. However, no software exists to analyze base editors, perform allele-specific quantification or that incorporates biologically-informed and scalable alignment approaches. Here, we present CRISPResso2 to fill this gap and illustrate its functionality by experimentally measuring and analyzing the editing properties of six genome editing agents.

---

The field of genome editing is rapidly advancing, and the technologies to modify the genome are becoming increasingly more accurate, efficient and versatile^1^. For example, base editors—a recent class of genome editing technology—harness the targeting properties of RNA-guided endonucleases to precisely change one nucleotide in a predictable manner^2,3,4^. As sequencing costs decrease and access to next-generation sequencing machines becomes more widespread, targeted amplicon sequencing is becoming the gold standard for the validation and characterization of genome editing experiments.

CRISPResso2 introduces five key innovations for the analysis of genome editing data: (1) Comprehensive analysis of sequencing data from base editors; (2) Allele specific quantification of heterozygous references; (3) A novel biologically-informed alignment algorithm; (4) Ultra-fast processing time; and (5) A batch mode for analyzing and comparing multiple editing experiments.

Existing software packages for the analysis of data generated by genome editing experiments are designed to only analyze cleavage events resulting from nuclease activity^5,6,7,8,9,10^. CRISPResso2 (http://crispresso2.pinellolab.org) is the first comprehensive software specifically designed to analyze base editor data from amplicon sequencing, in addition to quantifying and visualizing indels from other nucleases. CRISPResso2 allows users to readily quantify and visualize amplicon sequencing data from base editing experiments. It takes in raw FASTQ sequencing files as input and outputs reports describing frequencies and efficiencies of base editing activity, plots showing base substitutions across the entire amplicon region (**Fig. 1a**) and nucleotide substitution frequencies for a region specified by the user (**Fig. 1b**).

**Figure 1:**
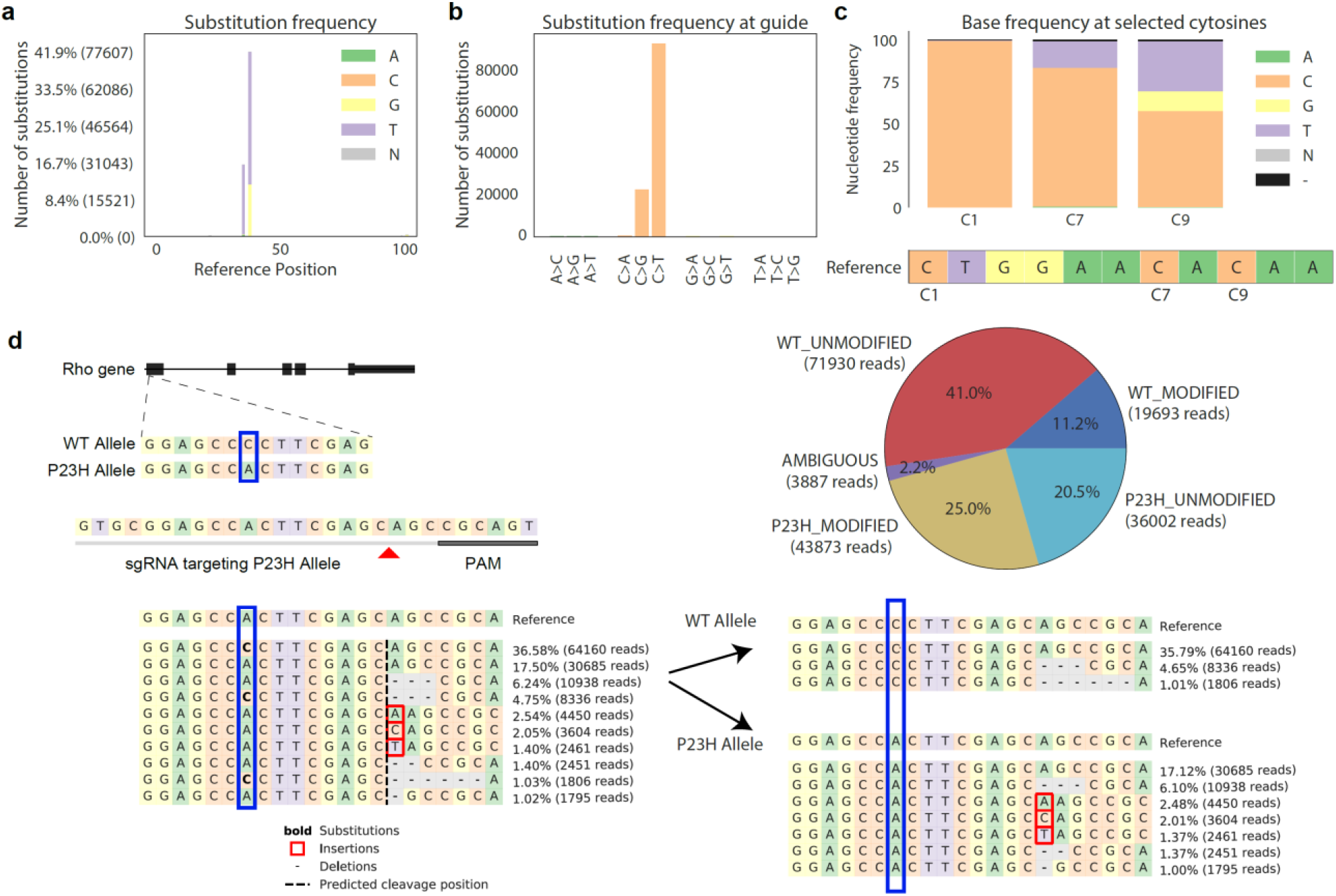
Novel features of CRISPResso2. a-c) CRISPResso2 analysis of base editing data. a) Locations of substitutions across the reference sequence. At each position, the number of substitutions to each non-reference base are shown. b) Barplot showing the frequency of substitution from a reference base to a non-reference base in the guide sequence. c) Percentage of each base at cytosines in the guide sequence. The reference sequence in the editing window is shown below the plot, with cytosines labeled according to position. For each cytosine, the frequency of each base at that position is shown. d) Allele-specific mutation calling for a guide targeting a mutated allele in the Rho gene in mice heterozygous for the mutated allele^11^. Reads (left) can be assigned to each allele using CRISPResso2 (right) to achieve accurate quantification of genome editing at genomic loci with multiple alleles. The pie chart shows the assignment of each read to the wild-type (red and dark blue) allele or to the P23H allele (yellow and light blue). Ambiguous alignments that could not be attributed uniquely to one of the alleles (e.g., due to a deletion at the SNP location) are shown in purple.

Additionally, users can specify the nucleotide substitution (e.g., C->T or A->G) that is relevant for the base editor used, and publication-quality plots are produced for nucleotides of interest with a heatmap showing conversion efficiency (**Fig. 1c**).

In cases where the genome editing target contains more than one allele, (for example when one or multiple heterozygous SNPs or indels are present), genome editing on each allele must be quantified separately, although reads from both alleles are amplified and mixed in the same input FASTQ file. Current strategies are not capable of analyzing multiple reference alleles and may lead to incorrect quantification. CRISPResso2 instead, enables allelic specific quantification by aligning individual reads to each allelic variant and assigning each read to the most closely-aligned allele. Downstream processing is performed separately for each allele so that insertions, deletions, or substitutions that distinguish each allele are not confounded with genome editing. To demonstrate the utility of our approach, we reanalyzed amplicon sequencing data from a mouse with a heterozygous SNP at the *Rho* gene where an engineered SaCas9-KKH nuclease was directed to the P23H mutant allele^11^. CRISPResso2 deconvoluted reads, quantified insertions and deletions from each allele, and produced intuitive visualizations of experimental outcomes (**Fig. 1d**).

Existing amplicon sequencing analysis toolkits ignore the biological understanding of genome editing enzymes and instead optimize the alignment based only on sequence identity. However, this can lead to incorrect quantification of indel events, especially in sequences with short repetitive subsequences where the location of indels may be ambiguous due to multiple alignments with the same best score. In such cases, it is reasonable to assume that indels should overlap with the predicted nuclease cleavage site. Our proposed alignment algorithm extends the Needleman-Wunsch algorithm with a mechanism whereby the assignment of insertions or deletions can be incentivized at specific indices in the reference amplicon sequence that correlate with predicted cleavage sites based on guide sequence and nuclease properties (**Supplementary note 1**). This increases the accuracy of indel calling and produces alignments that reflect our current understanding of the cleavage mechanism. We compared our improved alignment algorithm to those used in other amplicon-based genome editing analysis software and found that indels were incorrectly aligned to regions distal from the predicted cut site leading to incorrect quantification of editing events (**Supplementary note 2**).

As editing tools are refined and improved, and the possibility of therapeutic applications in humans approaches^1,12,13^, the importance of quantifying rare off-target mutagenesis has increased. In order to study putative off-targets, it is often necessary to analyze large-scale pooled sequencing datasets that profile hundreds of sites to assess the potential safety of genome editing interventions^14^. These and other large datasets have created a need for faster and more efficient analysis tools. To accelerate performance and decrease processing time, we designed and implemented an efficient implementation of our biologically-informed alignment algorithm. Further optimization of other components of the processing pipeline has reduced processing time ten-fold for large datasets, so that experiments analyzed using modern high-throughput sequencing technologies can be processed in under a minute as opposed several hours required by other software pacakges (**Supplementary Fig. 13**). We tested the accuracy of our new alignment algorithm and other optimizations using an extensive set of simulations with various mutational profiles and in the presence of sequencing errors and found that CRISPResso2 accurately recovered editing events with a negligible false-positive rate (<0.01) (**Supplementary note 3**).

Improvements in processing time and memory usage of CRISPResso2 have enabled users to analyze, visualize and compare results from hundreds of genome editing experiments using batch functionality of CRISPResso2. This is particularly useful when many input FASTQ files must to be aligned to the same amplicon or have the same guides, and the genome editing efficiencies and outcomes can be visualized together. In addition, we have created an intuitive plot that shows the nucleotide frequencies and indel rates at each position in each sample. This allows users to easily visualize the results and extent of editing in their experiments for different enzymes (Fig. 2a).

**Figure 2:**
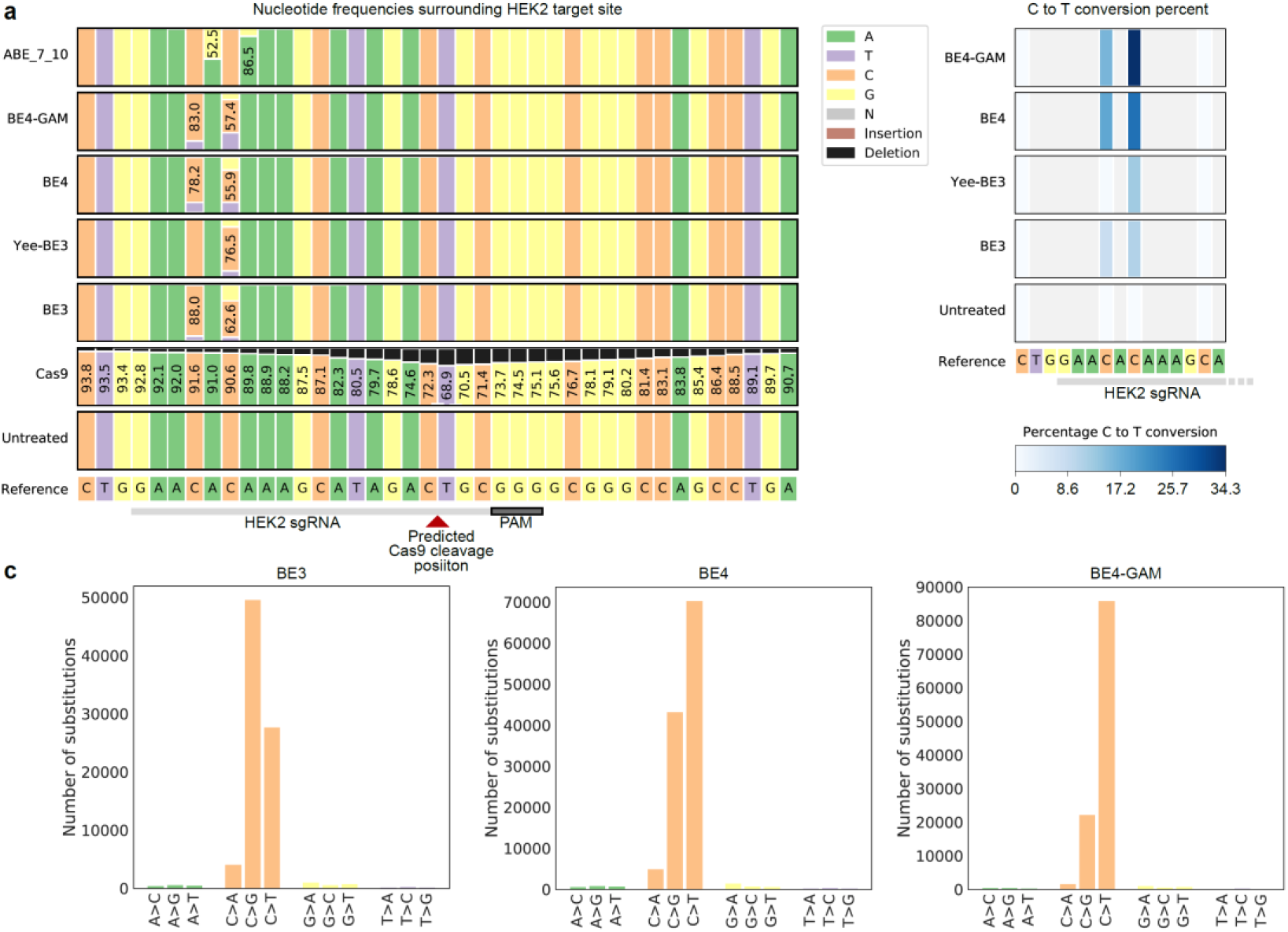
Batch analysis of genome editing data using CRISPResso2. *a) Batch output mode comparing six genome editing reagents and an untreated control. Each column represents the base composition at a single nucleotide at the HEK2 locus. The percentage of each base or indel at each nucleotide is shown as a proportion of each row. Bases with a frequency of between 50% and 99% are annotated with the percentage frequency of that nucleotide. b) Comparison of editing efficiencies of four base editors at the HEK2 protospacer sequence. The editing frequency at each cytosine is shown using the blue color scale, while non-cytosine bases are colored gray. c) Base editing purity for BE3, BE4, and BE4-GAM at the target site.*

Next, to showcase the utility and capabilities of CRIPSResso2 we generated a novel dataset to compare the mutational profiles of several modern genome editing tools. We performed genome editing via plasmid transfection in HEK293T cells using six genome editing agents: Cas9^15,16^, four C->T base editor variants (BE3^2^, Yee-BE3^3^, BE4^4^, BE4-GAM^4^) and one A->G base editor (Abe.7.10^17^) using guides targeting five well-studied genomic loci *(EMX1, FANCF,* and HEK293 sites 2, 3 and 4)^18^. Amplicon sequencing was performed at each target site for each guide-agent pairing and analyzed in batch mode by CRISPResso2 allowing us to easily analyze and compare the editing characteristics of each genome editing agent at each location (**Fig. 2a, Supplementary Fig. 2–5**).

Analysis of Cas9-treated samples at each guide target location suggests that the position of the most-frequently deleted bases varied between genomic loci, a finding which is similar to previous descriptions of guide-specific deletion profiles^19^. The HEK2, EMX1, and FANCF guides induced deletions at position 17 (from the 5’ end of the guide target), while Cas9-mediated deletion activity is predominantly observed at position 16 for the HEK3 and HEK4 guides (**Supplementary Fig. 6**). Insertions were predominantly observed between the bases at positions 17 and 18, although the frequencies of insertions varied between sites, with a maximum of 13.6% insertion frequency at the EMX1 locus and a minimum of 2.4% insertion frequency at the HEK4 locus (**Supplementary Fig.7**).

Likewise, we examined the base-editing preferences for each base editor and compared them across genomic locations. Here, we also observed guide-specific effects similar to those reported by others^20^, with most base editors showing the highest conversion rates at positions 4 to 9. However, at the C-rich FANCF site, all base editors showed extended conversion spectra even at the cytosine in the C_11_ position (Supplementary Fig.8–11).

We next examined patterns of genome editing at each site individually which can be easily visualized by CRISPResso2 in batch mode (**Fig. 2a**). Base editing replacement of C->T can be compared between each C->T base editor. For example, at the HEK2 locus (**Fig. 2b**), the editing preference of each base editor is noteworthy, with the preference for conversion of the cytosine at the C_6_ position, as compared to the cytosine at the C4 position. Notably, the Yee-BE3 editor has a shifted activity window resulting in very little base editing activity at the C4 position (**Supplementary Fig. 12**).

Overall editing efficiencies for each site can also be visualized using detailed plots (**Fig. 2c**). BE4-GAM has the highest C->T purity, meaning that most editing products involve C->T changes. In contrast, BE3 creates more C->G changes at the HEK2 site. The increased product purity of BE4-GAM can be attributed to the addition of Gam protein from bacteriophage Mu which reduces indel formation during base editing^4^.

In summary, CRISPResso2 is a software tool for the comprehensive analysis, visualization and comparison of sequencing data from genome editing experiments. In addition to accurate indel analysis from nucleases such as Cas9, CRISPResso2 offers analysis tools for recent base editors, support for multiple alleles, increased computational speed, an improved alignment algorithm, and a batch functionality for analyzing and comparing genome editing experiments across hundreds of samples. CRISPResso2 is available online at http://crispresso2.pinellolab.org.

## Online Methods

### Cell culture

HEK293T (American Type Culture Collection (ATCC) CRL-3216) were maintained in DMEM plus GlutaMax (Thermo Fisher) supplemented with 10% (v/v) fetal bovine serum, at 37 °C with 5% CO_2_. Cells were verified to be mycoplasma-free upon purchase, and mycoplasma testing was performed every 6 months to ensure cells were mycoplasma free.

### Plasmid transfection and cell harvest

HEK293T cells were seeded one day prior to transfection at a density of 30,000 cells per well on 48-well collagen-coated BioCoat plates (Corning).

Plasmids were midi-prepped for transfection using the ZymoPURE II Plasmid Kit (Zymo Research) according to the manufacturer’s instructions. 750 ng of base editor or Cas9 plasmid and 250 ng of sgRNA expression plasmid was transfected using 1.5 ml of Lipofectamine 2000 (Thermo Fisher) per well according to the manufacturer’s protocol. 3 days post-transfection, cells were harvested in lysis buffer (10 mM Tris-HCl, pH 7.0, 0.05% SDS, 25 μg/mL Proteinase K (Thermo Fisher Scientific)) and incubated at 37 °C for 60 mins, followed by 20 mins at 80 °C to denature Proteinase K. Genomic DNA was amplified by PCR and subjected to high throughput sequencing using an Illumina MiSeq according to previously published protocol^20^.

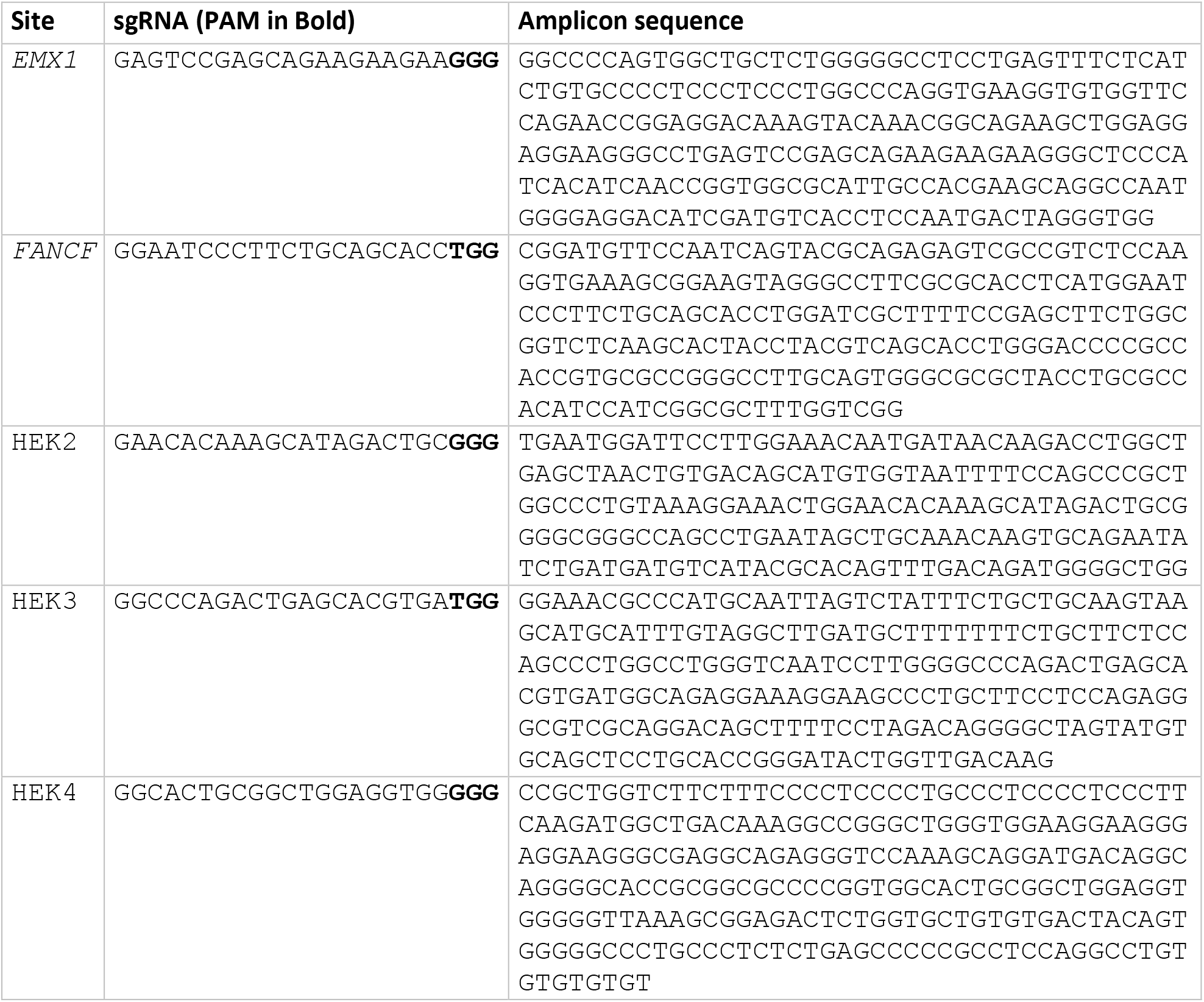
Guide RNAs and amplicon sequences used

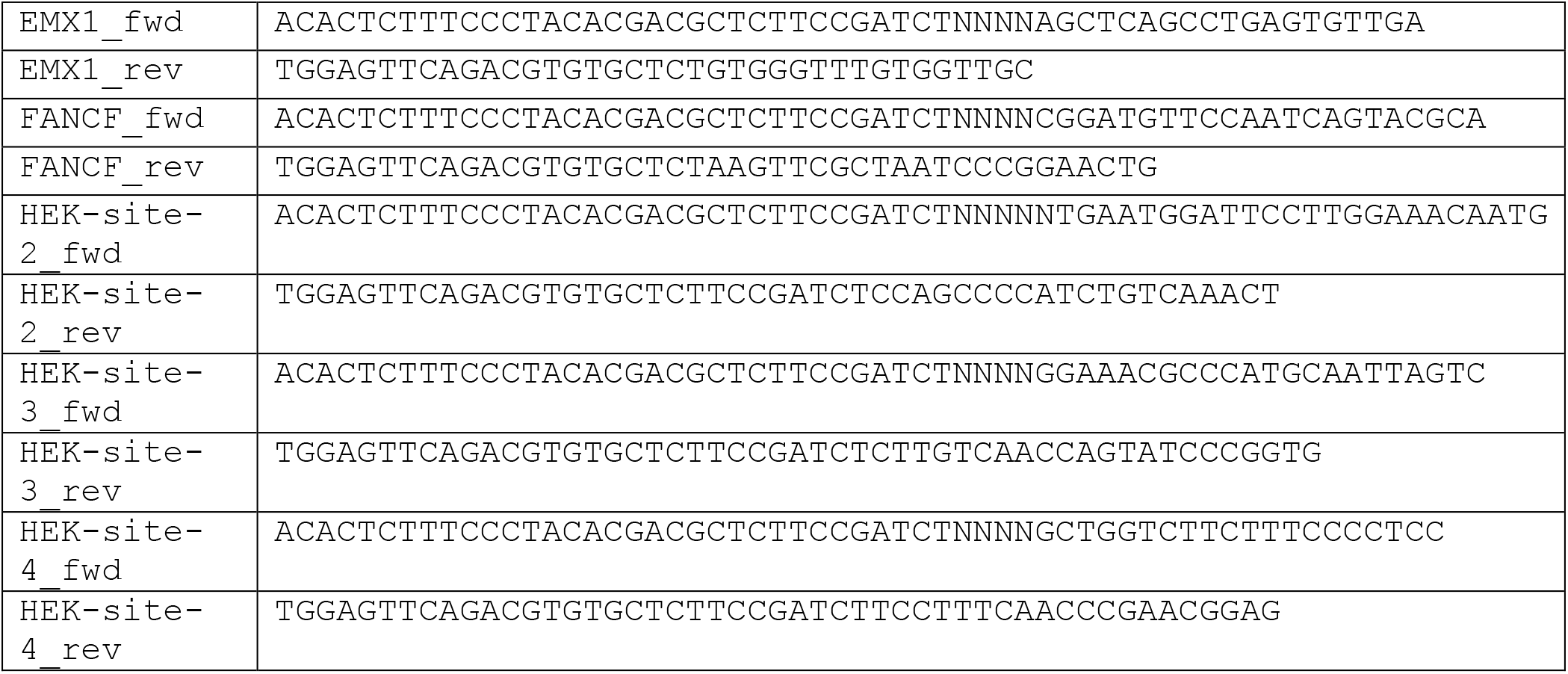
Sequencing primers used

### Data Analysis

Next-generation sequences were analyzed using CRISPResso2.

## Acknowledgements

D.R.L. is supported by DARPA HR0011-17-2-0049; U.S. NIH RM1 HG009490, R01 EB022376, and R35 GM118062; and HHMI. D.E.B. is supported by NIDDK (R03DK109232), NHLBI (DP2OD022716, P01HL32262), Burroughs Wellcome Fund, Doris Duke Charitable Foundation, and ASH Scholar Award. L.P. is supported by the National Science Foundation National Institutes of Health R00HG008399 and the Defense Advanced Research Projects Agency HR0011-17-2-0042.

## Author contributions

K.C. and L.P. conceived the project, led the study, and wrote the software. H.R. with input from D.R.L. performed biological experiments. K.C. analyzed experimental data. All authors contributed input on measurement and visualization of genome editing outcomes and provided input on the manuscript.

## Supplementary Figures

**Supplementary Figure 1:**
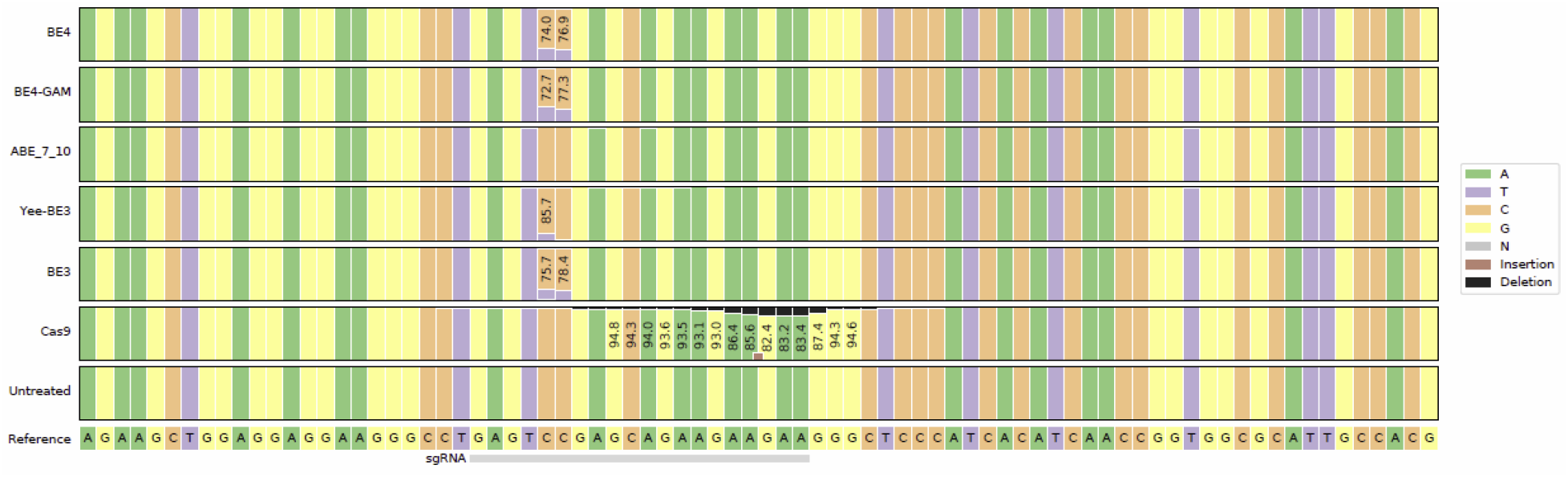
CRISPResso2 output from genome editing at *EMX1*

**Supplementary Figure 2:**
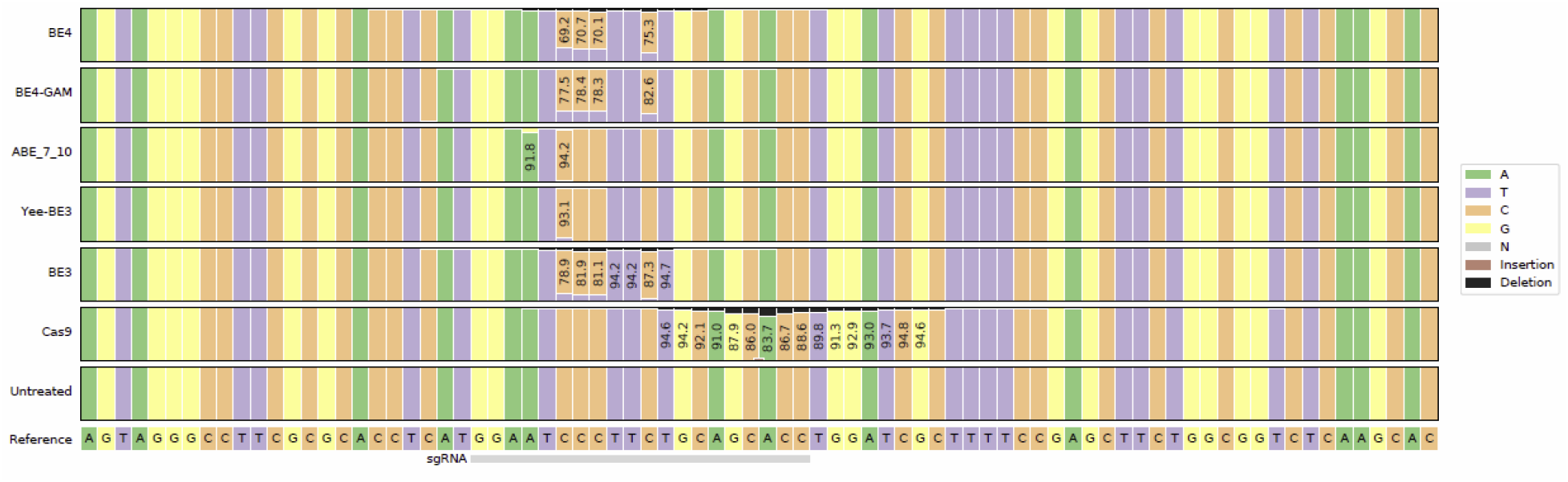
CRISPResso2 output from genome editing at *FANCF*

**Supplementary Figure 3:**
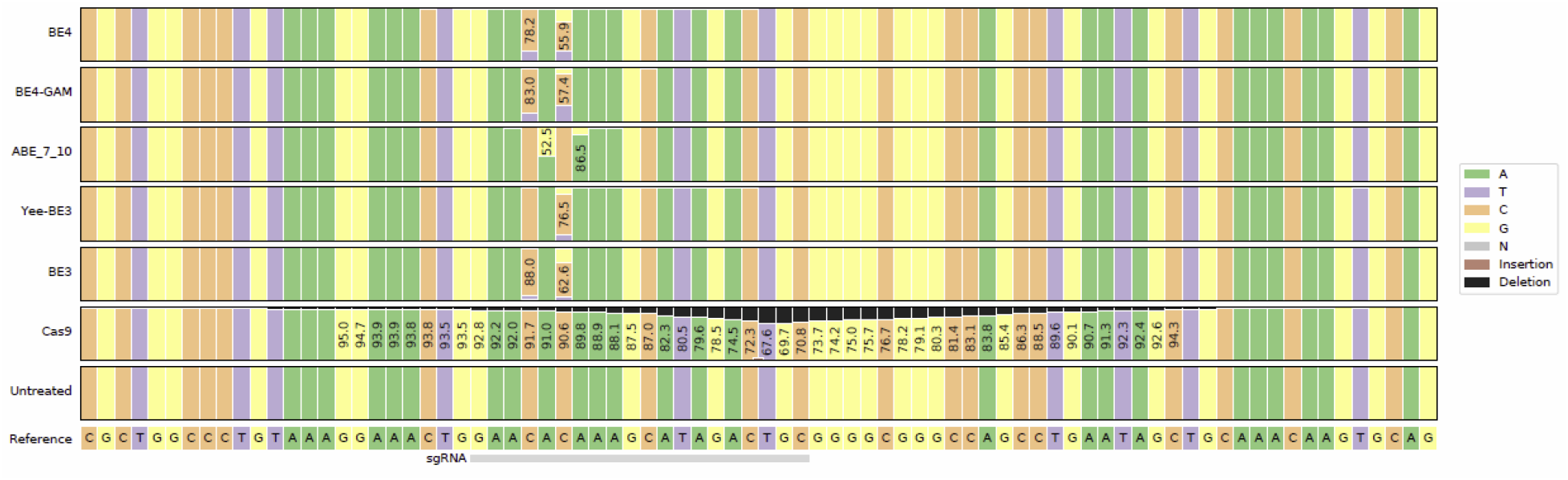
CRISPResso2 output from genome editing at HEK2

**Supplementary Figure 4:**
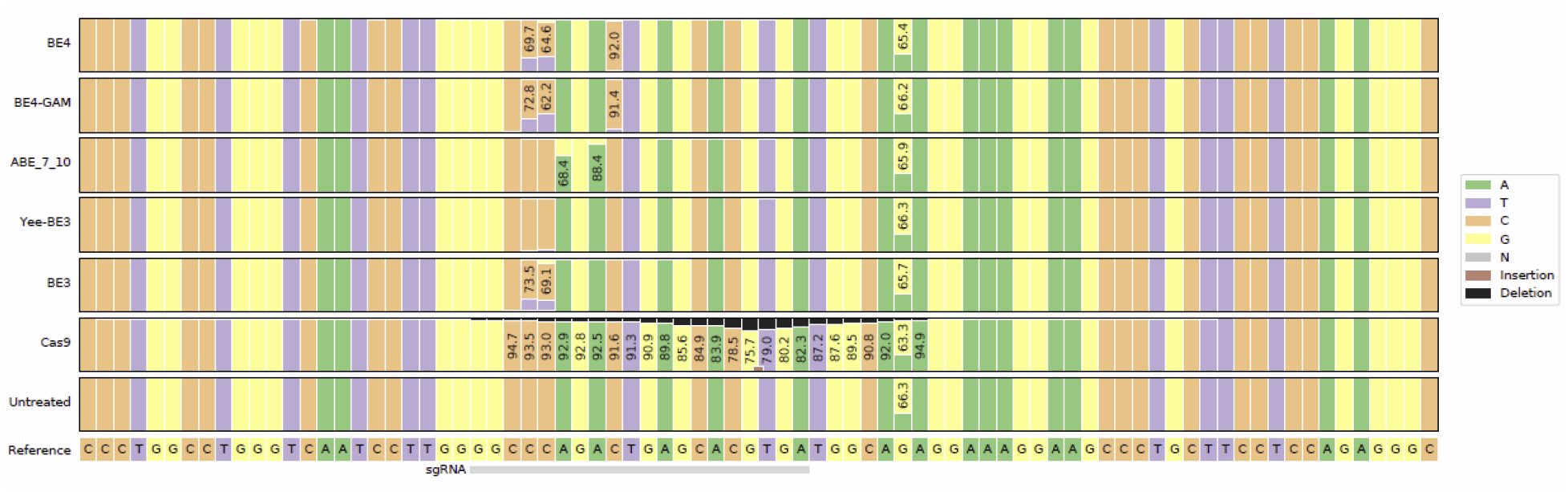
CRISPResso2 output from genome editing at HEK3

**Supplementary Figure 5:**
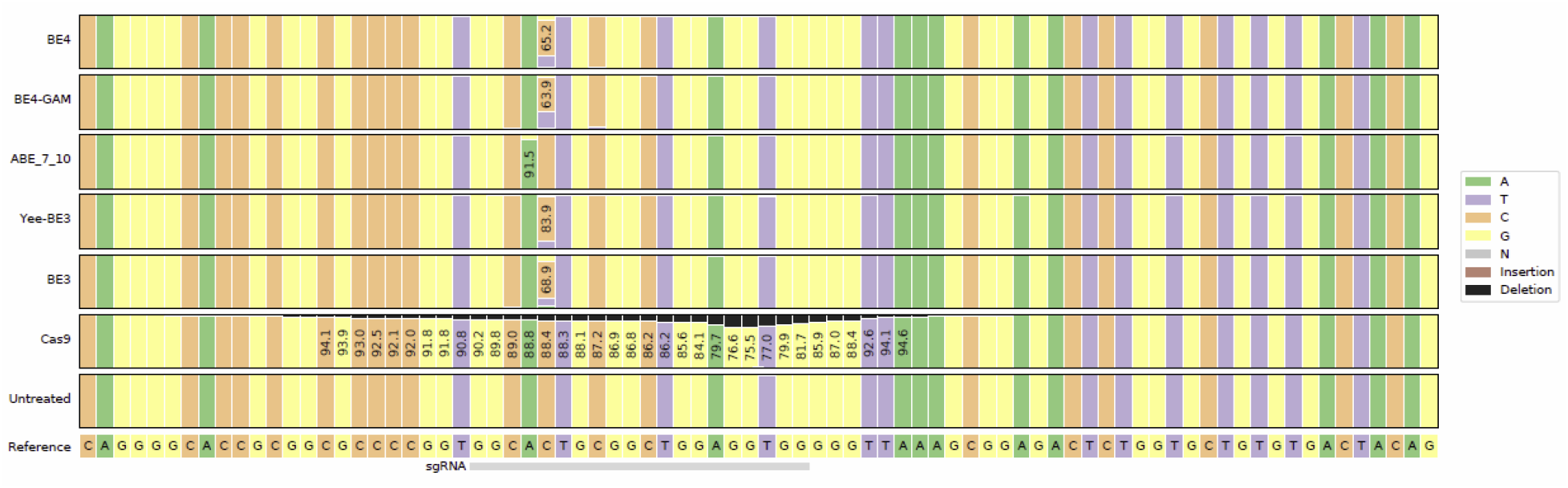
CRISPResso2 output from genome editing at HEK4

**Supplementary Figure 6:**
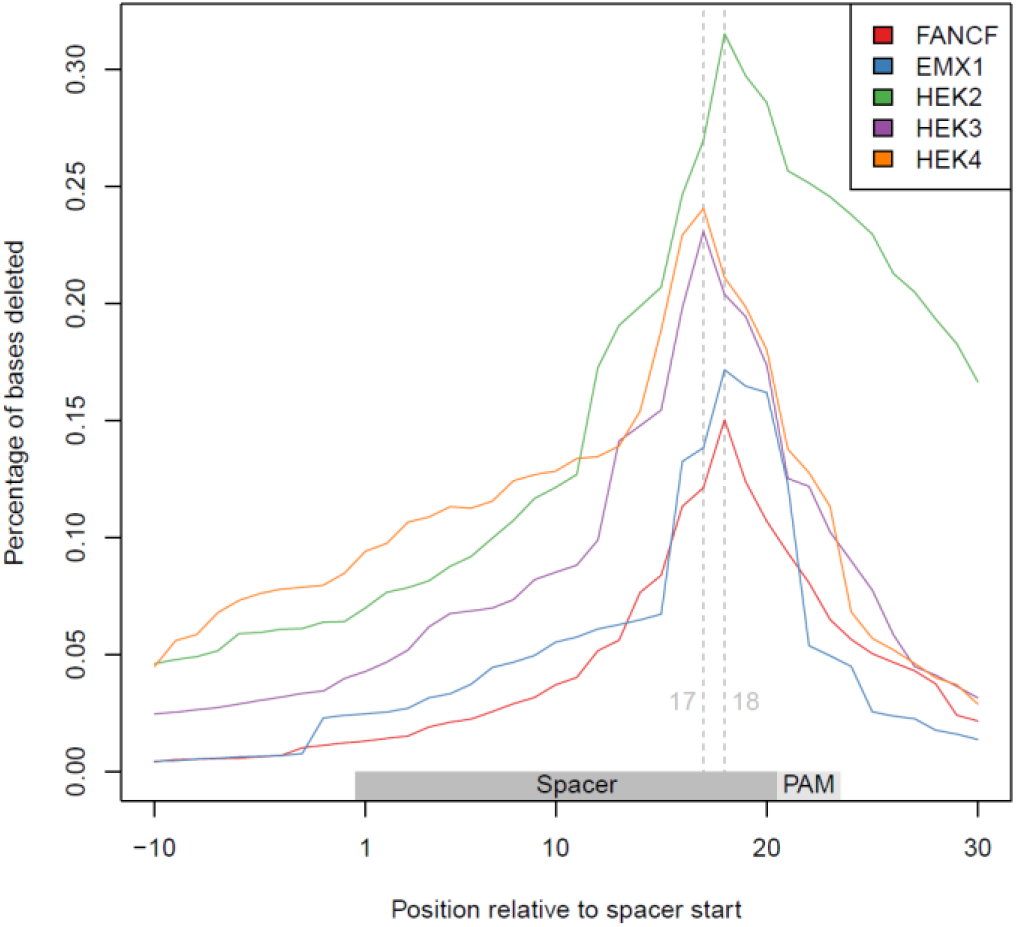
Location of deletion events across Cas9 samples. Dashed vertical lines mark nucleotides positions 17 and 18 in reference to the start of the sgRNA spacer sequence.

**Supplementary Figure 7:**
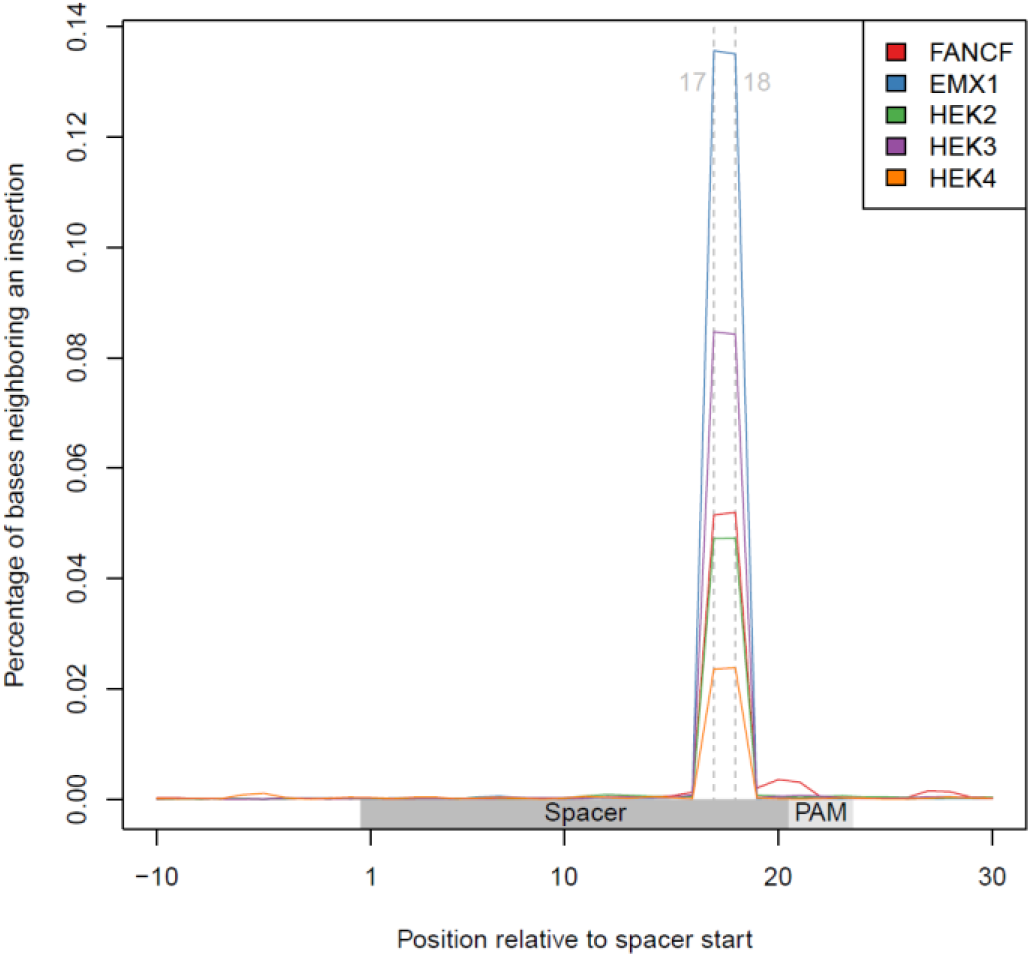
Location of insertion events across Cas9 samples. Dashed vertical lines mark nucleotides positions 17 and 18 in reference to the start of the sgRNA spacer sequence.

**Supplementary Figure 8:**
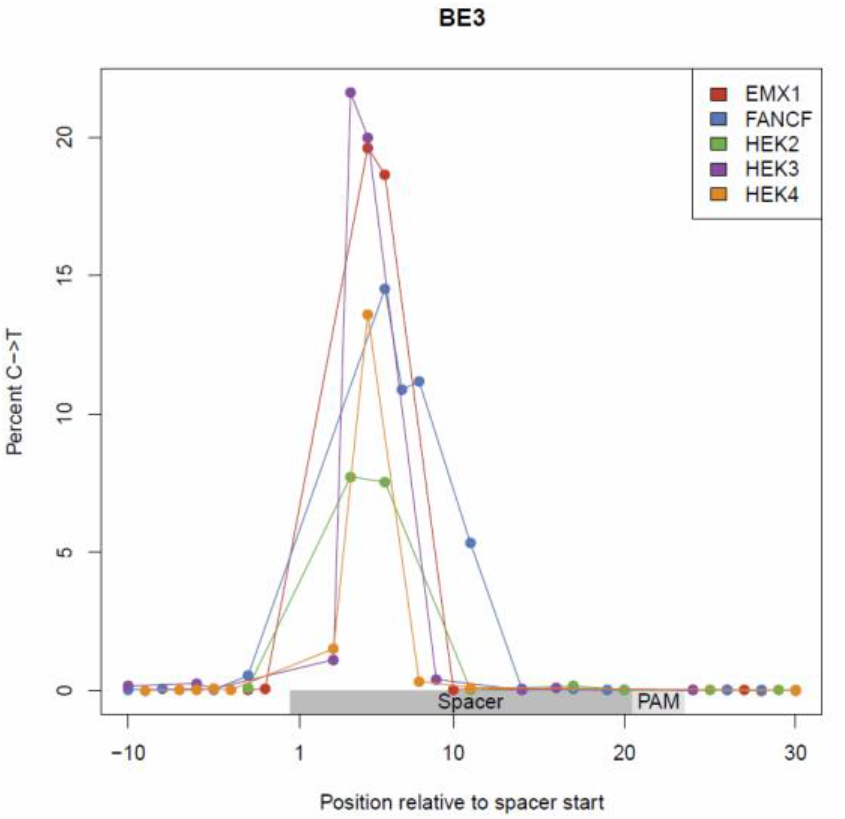
Base editing (C->T) percentage in samples treated with BE3. BE3

**Supplementary Figure 9:**
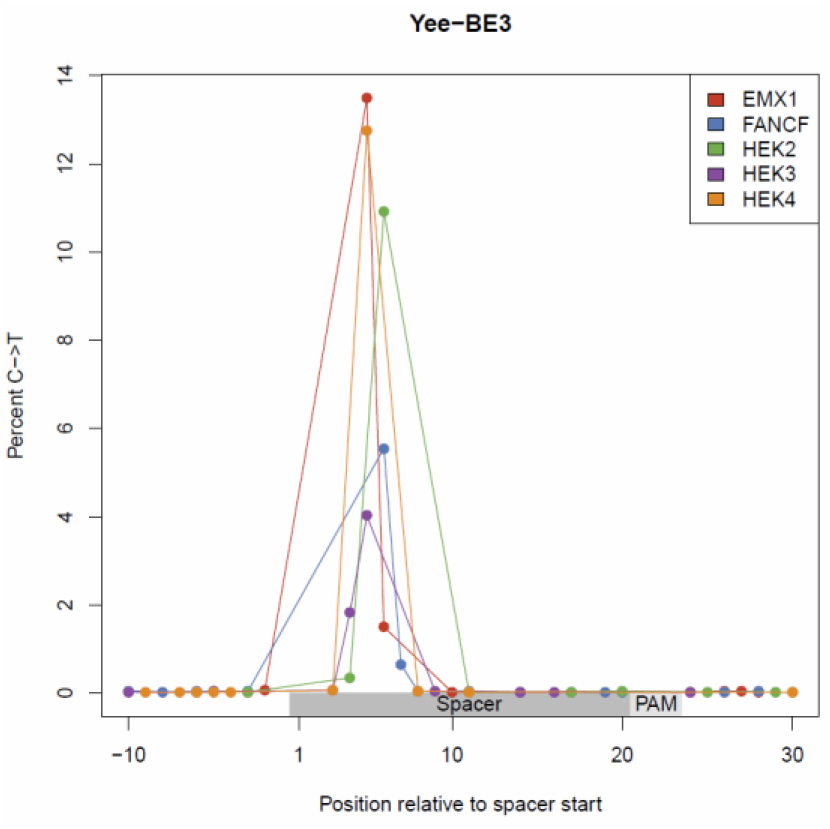
Base editing (C->T) percentage in samples treated with Yee-BE3.

**Supplementary Figure 10:**
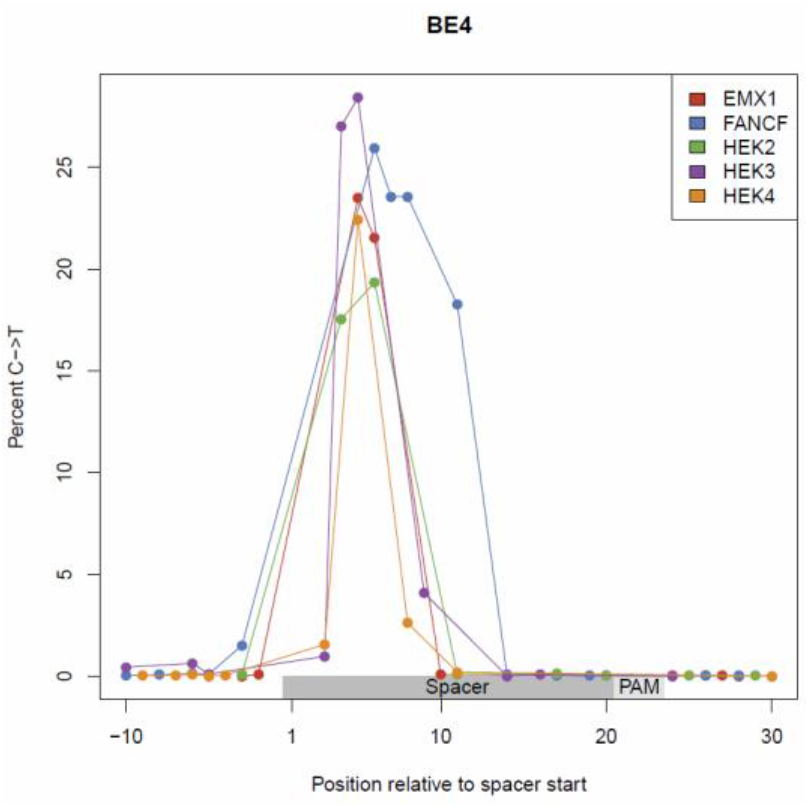
Supplementary Figure 10: Base editing (C->T) percentage in samples treated with BE4. BE4

**Supplementary Figure 11:**
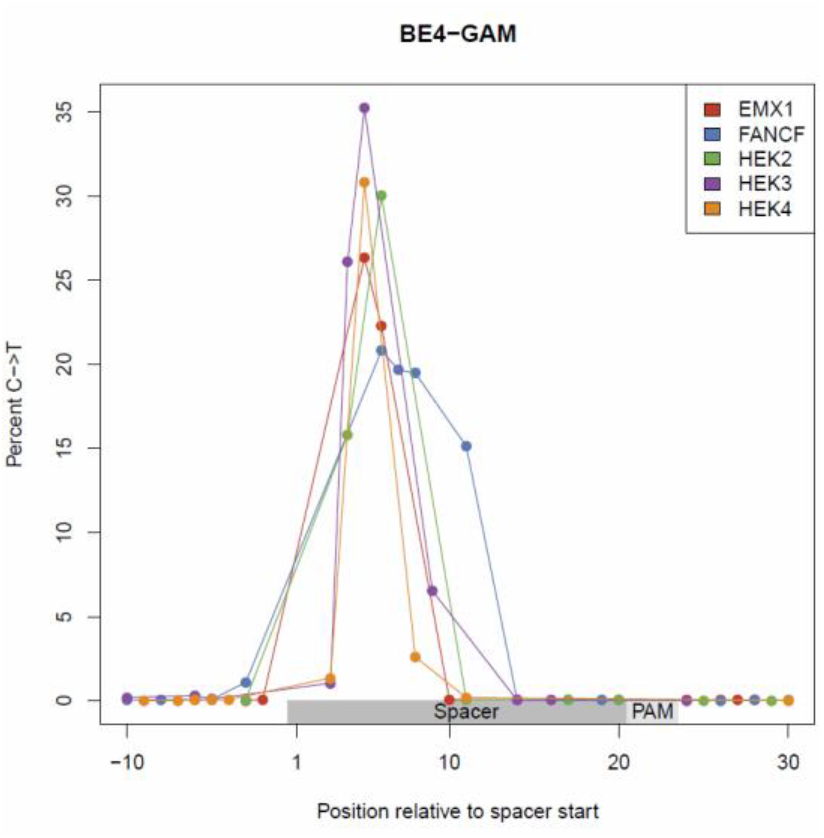
Base editing (C->T) percentage in samples treated with BE4-GAM.

**Supplementary Figure 12:**
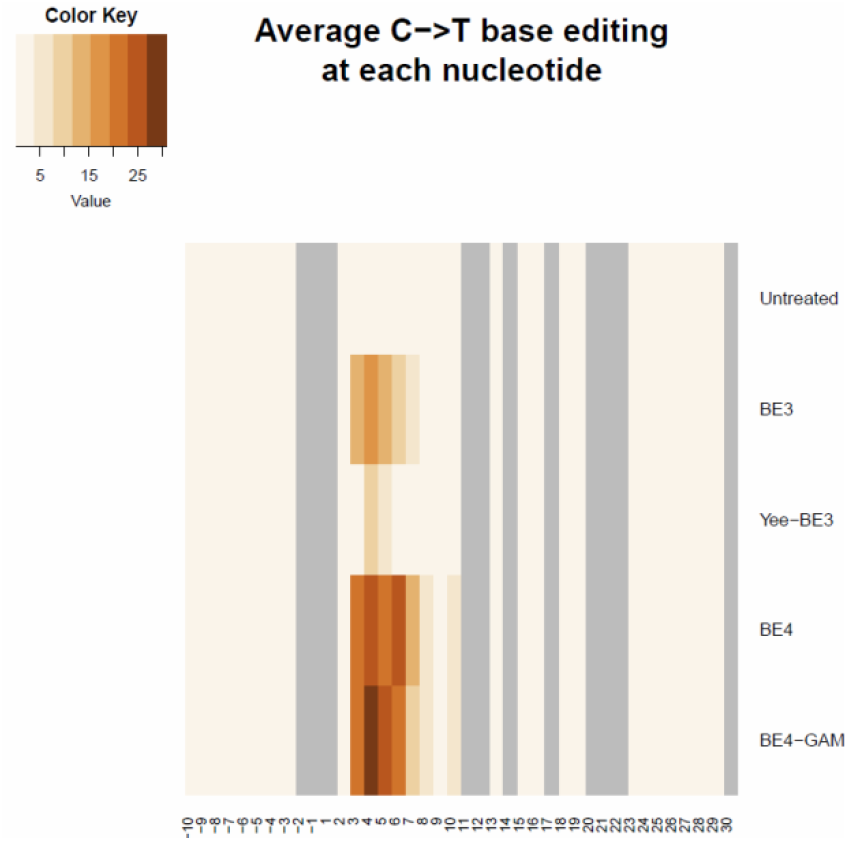
Average base editing (C->T) percentage across all guides

**Supplementary Figure 13:**
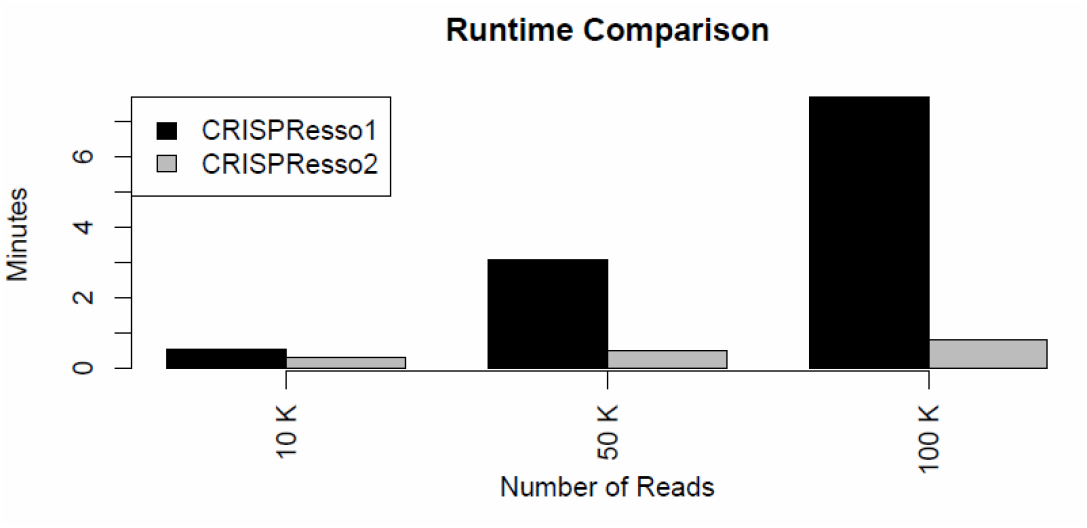
Runtime comparison between CRISPResso and CRISPResso2 Runtime Comparison

## Supplementary Notes

Supplementary Note 1: Details on alignment algorithm improvements

Existing alignment algorithms find an optimal alignment score based on a match score, and penalties for gaps. An affine gap penalty structure allows consecutive gaps to be penalized with a ‘gap extension’ score which is generally lower than the ‘gap open’ score. This tends to favor alignments with clusters of insertions or deletions reflecting the biological intuition that insertions and deletions are caused by a single nuclease cleavage event rather than multiple cleavages that create multiple single-base deletions or insertions.

For many alignments between a reference sequence and a sequencing read, one single alignment produces a highest alignment score, and that is assumed to be the correct alignment. However, particularly if the sequence is repetitive around the putative insertion or deletion, multiple alignments will share the same best score – meaning that the assignment of insertions and deletions across the reference sequence can occur at multiple locations and return the same score. Our understanding of nuclease activity suggests that most indels occur around a predicted cut site which is a physical characteristic of each nuclease. In order to select alignments with indels at the predicted cut site, we implemented a modified version of the Needleman-Wunsch (Needleman 1970) or Gotoh (Gotoh 1990) alignment that incentivizes indels at the predicted cut site while using an affine gap penalty scoring metric.

Given

- The match/mismatch score between nucleotide at position *i* and *j* as *S_ij_*
- Gap opening penalty as *GO*
- Gap extension penalty as *GE*
- the gap incentive at position *i* as *GI_i_*

The objective is to find the optimal alignment between read *A* and the reference sequence *B*. The matrices *I, J*, and *M* (each with dimensions length(A)xlength(B)) can be filled using dynamic programming. *M_ij_* represents the score of the best alignment between *A[0,i]* (meaning the first *i* characters of A) and *B[0,j]* that ends with a match between *A_i_* and *B_j_*. *I_i,j_*, represents the score of the best alignment between *A[0,i]* and *B[0,j]* that ends with a gap being paired with *A_i_. J_i,j_* represents the score of the best alignment between *A[0,i]* and *B[0,j]* that ends with a gap being paired with *B_j_*. After all three matrices are computed, a traceback procedure is used to recover the optimal aligment.

The Gotoh algorithm defines the score at each position in matrices I, J, and *M* recursively as:

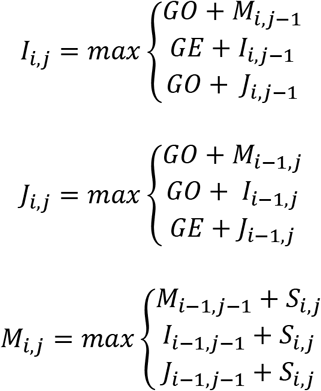

Our modification to the algorithm incorporates the gap incentive vector GI, and also removes the possibility of following a gap in one sequence with a gap in the next sequence, which is the biological equivalent of observing an insertion and a deletion at the same basepair. These two modifications are reflected in our definitions below:

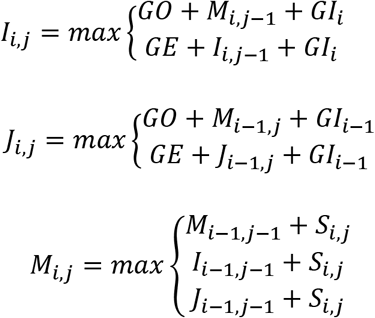

Supplementary Note 2: Comparison of alignment strategies from other amplicon-based genome editing software The alignment of insertions and deletions can have an important effect on quantification of genome editing. We demonstrate this with a simulated reference and read produced by a 2bp insertion at the predicted Cas9 cut site.

Reference sequence:

GGGTGGGCTACAAGAGTGCAAGCTATCAGTTGCTTCTATATCACAGCCTCGACGAATGGTATGGCCTGTACCAGGGTCAATA

Read Sequence:

GGGTGGGCTACAAGAGTGCAAGCTATCAGTTGCTTCTATATATCACAGCCTCGACGAATGGTATGGCCTGTACCAGGGTCAATA

Guide Sequence:

ATCAGTTGCTTCTATATCAC

We created a simulated fastq with five copies of the read sequence for testing analysis on different CRISPR analysis toolkits.

@M0000:100:000000000-ABCD:1:0001:0001:0001 1:N:0:CGAT

GGGTGGGCTACAAGAGTGCAAGCTATCAGTTGCTTCTATATATCACAGCCTCGACGAATGGTATGGCCTGTACCAGGGTCAATA +

CCCCCCCFFFFF5GGGGGGGGGHHHHHHHGGGGGGHHHHHHHHHHHDGGGGEEGHGHHGGGGGGHHHHHHGHHHHHHHGGGGGH @M0000:100:000000000-ABCD:1:0001:0001:0002 1:N:0:CGAT

GGGTGGGCTACAAGAGTGCAAGCTATCAGTTGCTTCTATATATCACAGCCTCGACGAATGGTATGGCCTGTACCAGGGTCAATA +

CCCCCCCFFFFF5GGGGGGGGGHHHHHHHGGGGGGHHHHHHHHHHHDGGGGEEGHGHHGGGGGGHHHHHHGHHHHHHHGGGGGH @M0000:100:000000000-ABCD:1:0001:0001:0003 1:N:0:CGAT

GGGTGGGCTACAAGAGTGCAAGCTATCAGTTGCTTCTATATATCACAGCCTCGACGAATGGTATGGCCTGTACCAGGGTCAATA +

CCCCCCCFFFFF5GGGGGGGGGHHHHHHHGGGGGGHHHHHHHHHHHDGGGGEEGHGHHGGGGGGHHHHHHGHHHHHHHGGGGGH @M0000:100:000000000-ABCD:1:0001:0001:0004 1:N:0:CGAT

GGGTGGGCTACAAGAGTGCAAGCTATCAGTTGCTTCTATATATCACAGCCTCGACGAATGGTATGGCCTGTACCAGGGTCAATA +

CCCCCCCFFFFF5GGGGGGGGGHHHHHHHGGGGGGHHHHHHHHHHHDGGGGEEGHGHHGGGGGGHHHHHHGHHHHHHHGGGGGH @M0000:100:000000000-ABCD:1:0001:0001:0005 1:N:0:CGAT

GGGTGGGCTACAAGAGTGCAAGCTATCAGTTGCTTCTATATATCACAGCCTCGACGAATGGTATGGCCTGTACCAGGGTCAATA +

CCCCCCCFFFFF5GGGGGGGGGHHHHHHHGGGGGGHHHHHHHHHHHDGGGGEEGHGHHGGGGGGHHHHHHGHHHHHHHGGGGGH

We performed analysis on this fastq with the given amplicon reference and guide sequence using CRISPResso2 which produced the following alignment:

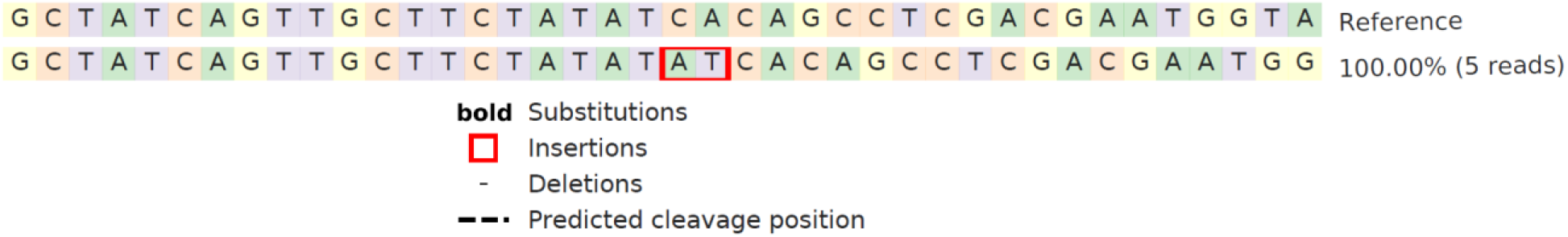

Note that the insertion is correctly placed at the 3bp away from the PAM, at the predicted cleavage position, and that all reads are identified as ‘modified’ reads.

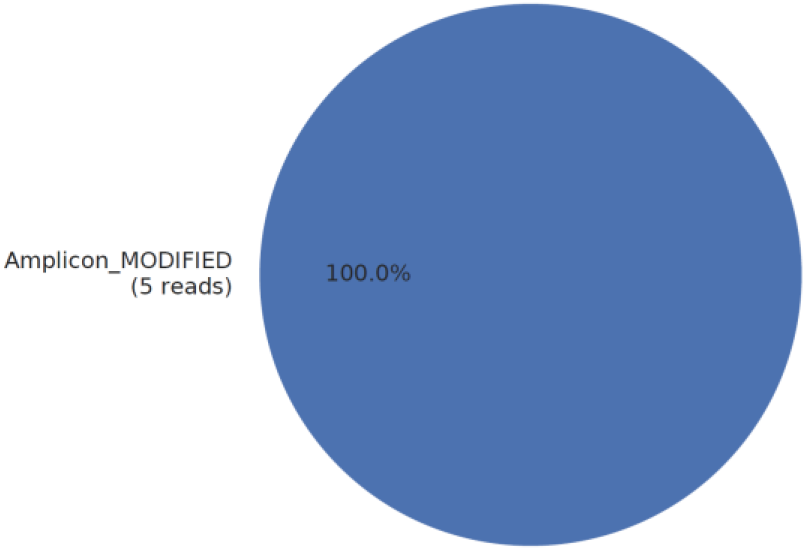

Alignments that are not aware of the predicted cutting at position 17 will produce alternate alignments.

**Cas-Analyzer** (Park 2017) aligns the indel to the left of the predicted cut site and not at the predicted cut site.

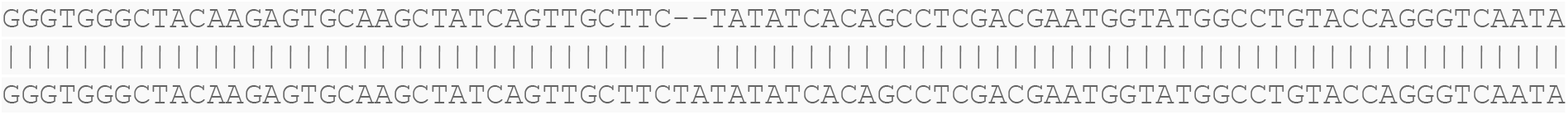

Cas-Analyzer excludes spurious indels that may be due to factors other than nuclease cleavage by only considering indel events that occur within a certain window (default 5bp) of the predicted cut site. Because the indel was aligned incorrectly, the indel is excluded from analysis.

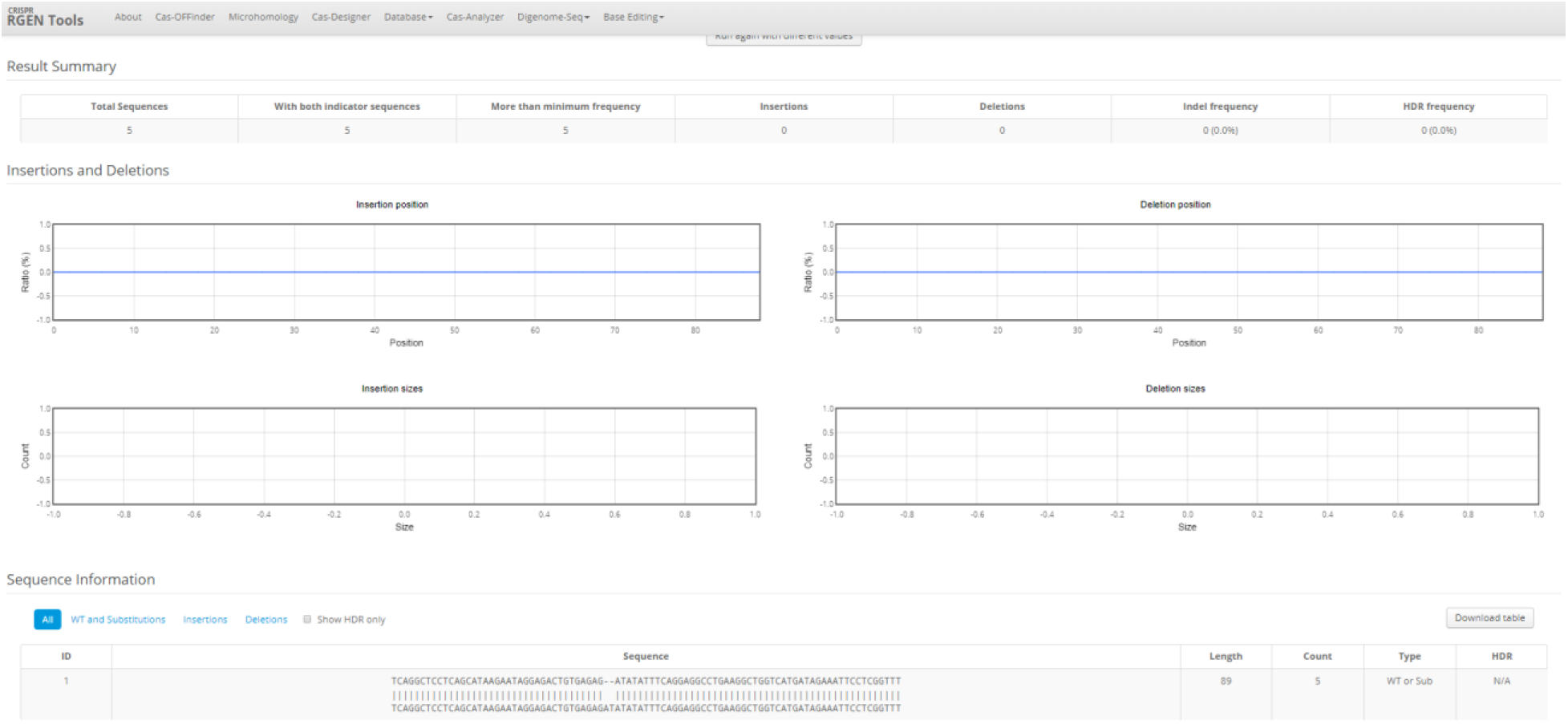

Note that the two basepair insertion has been detected, but because it has been aligned outside of the predicted cut site, the insertion is not counted, and all reads are called unedited:

| Total Sequences | With both indicator sequences | More than minimum frequency | Insertions | Deletions | Indel frequency | HDR frequency |
| --- | --- | --- | --- | --- | --- | --- |
| 5 | 5 | 5 | 0 | 0 | 0 (0.0%) | 0 (0.0%) |

CRISPR-DAV (Wang 2017) uses BWA (Li 2009) and ABRA (Mose 2014) to align reads to a reference. The resulting alignments assign insertion away from the predicted cleavage site:

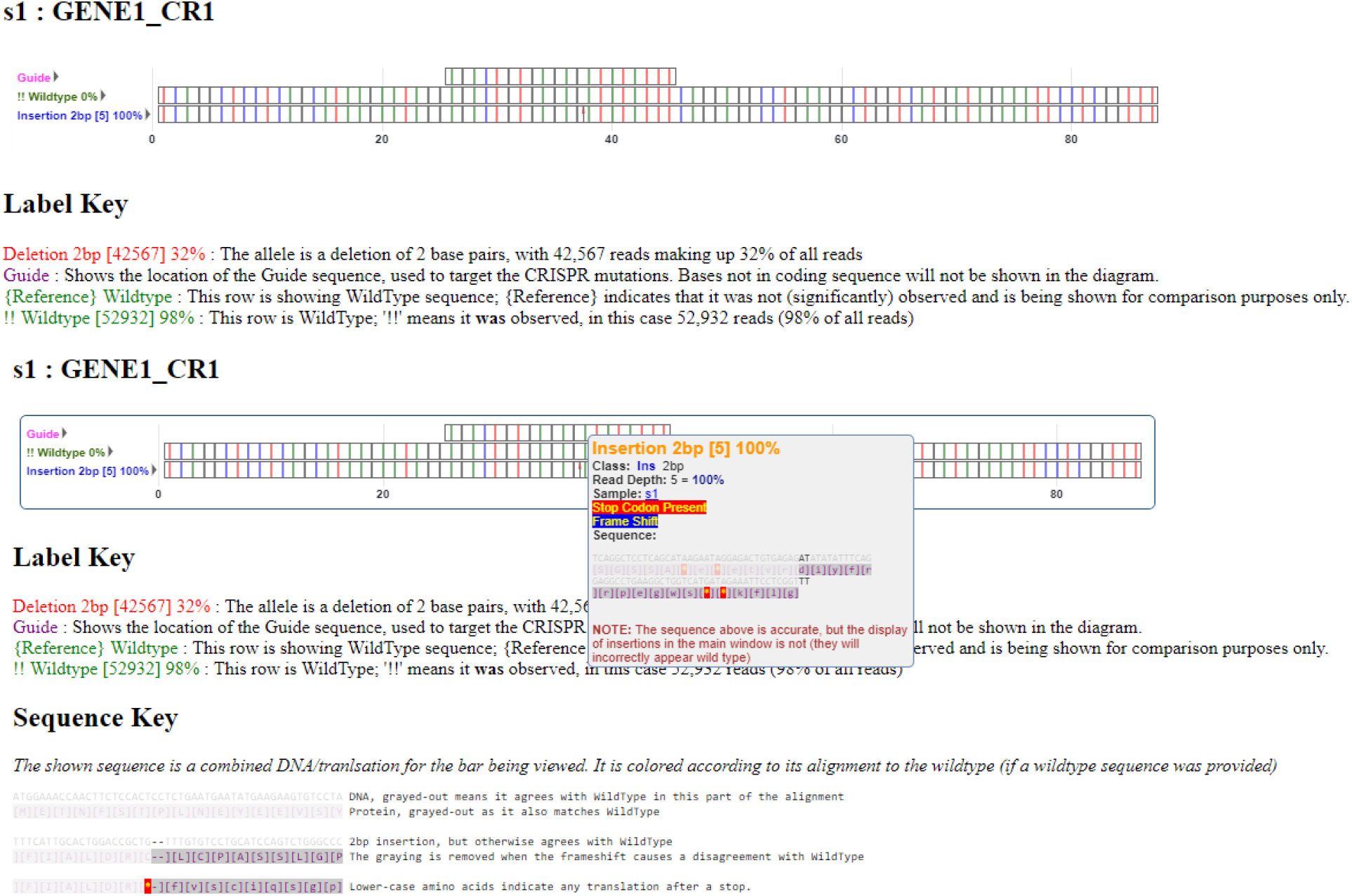

---

Supplementary note 3: Measurement of calling accuracy by simulation studies

To measure the accuracy of indel calling and the robustness against sequencing errors, we simulated a several datsets and measured the ability of CRISPResso2 to recover indels.

We first simulated 10 datasets with no sequencing error with 1000 unmodified reads. To 9 of the datasets, we added 1000 reads with the following modifications: substitutions (1,2,3bp), deletions (5,10,50bp), and insertions (5,10,50bp). We measured the number of modified reads called by CRISPResso2.

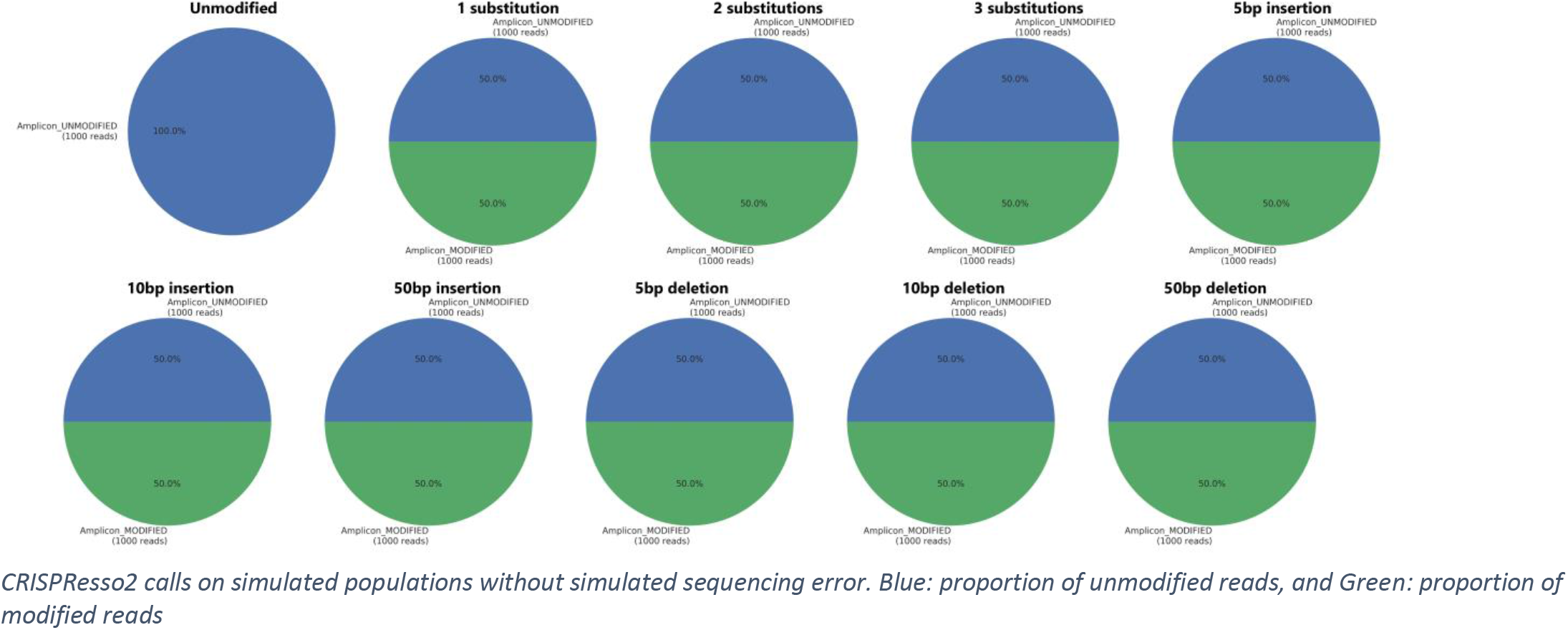

We then resimulated the datasets but introduced sequencing errors similar to those produced by the Illumina Miseq platform using the ART tool (Huang 2012). We used CRISPResso2 to quantify the number of modified and unmodified alleles.

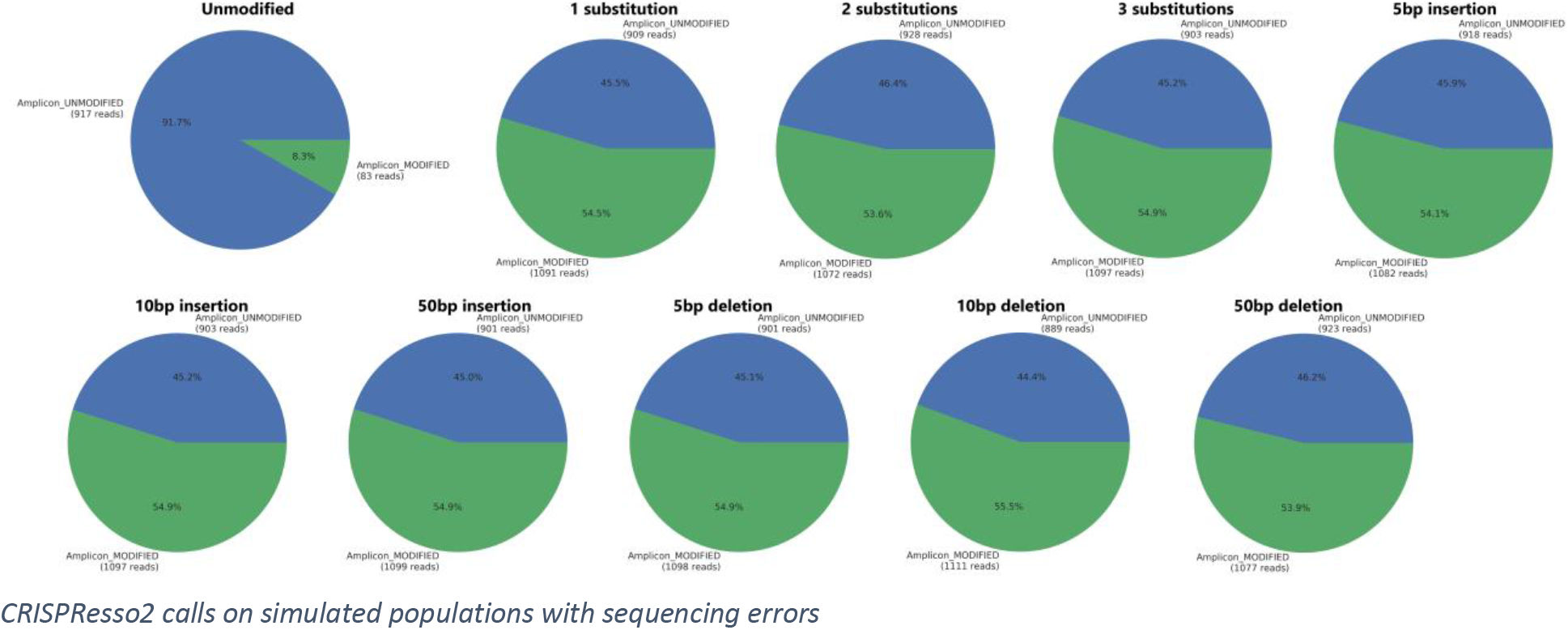

We then ran CRISPResso2 on the simulated reads with simulated sequencing errors but set 2bp window around the predicted cleavage site (1bp on each side) ignoring modifications outside of this window.

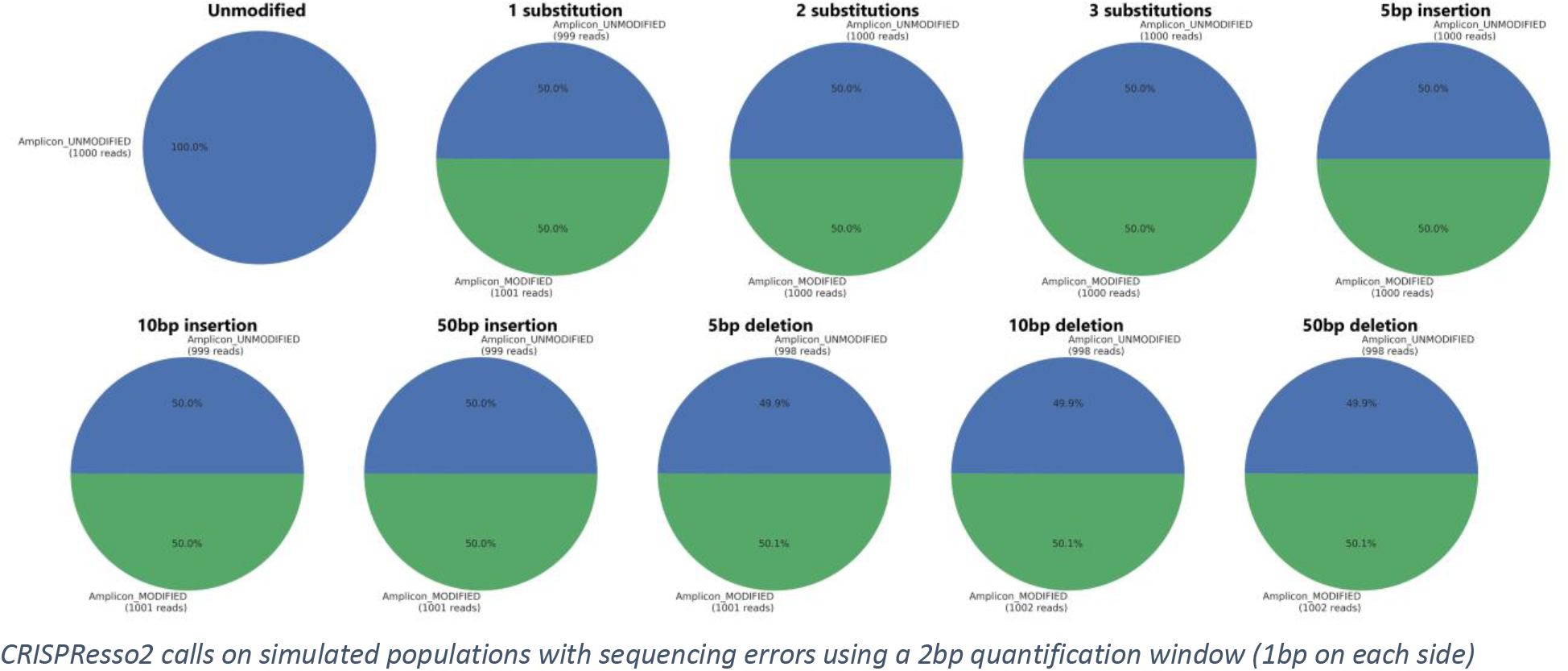

These three results show that by using the quantification window of CRISPResso2, accurate rates of allele modification can be recovered even in the presence of sequencing errors.

